# The RNA Virome of Echinoderms

**DOI:** 10.1101/2022.03.01.482561

**Authors:** Elliot W. Jackson, Roland C. Wilhelm, Daniel H. Buckley, Ian Hewson

## Abstract

Echinoderms are a phylum of marine invertebrates that include model organisms, keystone species, and animals commercially harvested for seafood. Despite their scientific, ecological, and economic importance, there is little known about the diversity of RNA viruses that infect echinoderms compared to other invertebrates. We screened over 900 transcriptomes and viral metagenomes to characterize the RNA virome of 38 echinoderm species from all five classes (Crinoidea, Holothuroidea, Asteroidea, Ophiuroidea and Echinoidea). We identified 347 viral genome fragments that were classified to genera and families within nine viral orders - *Picornavirales, Durnavirales, Martellivirales, Nodamuvirales, Reovirales, Amarillovirales, Ghabrivirales, Mononegavirales*, and *Hepelivirales*. We compared the relative viral representation across three life stages (embryo, larvae, adult) and characterized the gene content of contigs which encoded complete or near-complete genomes. The proportion of viral reads in a given transcriptome was not found to significantly differ between life stages though the majority of viral contigs were discovered from transcriptomes of adult tissue. This study illuminates the biodiversity of RNA viruses from echinoderms, revealing the occurrence of viral groups in natural populations.

## Introduction

Metazoans harbor an enormous diversity and abundance of RNA viruses – a discovery that has reshaped our understanding of viral evolution through expanded viral-host associations, broadened phylogenetic diversity, and novel reconfigurations of genome architectures [1, 2]. Newly discovered viruses often blur the boundaries between well-known viral groups. For example, prior to a recent expansion, the family *Flaviviridae* was typified by relatively uniform, monopartite genomes, having a single 10-12 kb-long open reading frame (ORF), and being vectored to mammals by arthropod. Metagenomics has led to the discovery of hundreds of novel flavivirus genomes, redefining the genomic properties of this viral family, and extending their host diversity beyond mammals [3–6]. To date, the exploration and systematization of invertebrate RNA viruses have been skewed towards terrestrial arthropods, mainly insects, leaving gaps in our understanding of the diversity, ecology, and evolution of RNA virus in other invertebrate groups [1, 7–12]. To help close this gap, we characterized the RNA virome of Echinodermata - a phylum of marine invertebrates that are globally distributed throughout Earth’s oceans and represent an evolutionary crossroads in developmental biology as one of two phyla that are invertebrate deuterostomes.

Wildlife disease and aquaculture are the two primary areas of concern regarding the threat of viral outbreaks among echinoderms. Disease outbreaks of sea urchins and sea stars have been documented at local, regional, and continental scales since 1898 and have gone unresolved in regards to their etiology [13, 14]. Certain species of sea urchins and sea cucumbers are valued as seafood delicacies and the growing demands for these species in the seafood industry have led to a rise in aquaculture farming [15, 16]. Viral outbreaks pose a major concern for aquaculture operations [17, 18], yet little is known about the identity, let alone virulence, of viruses that infect these animals [19–21]. Baseline knowledge of viruses present in wild populations can help determine the etiology of future outbreaks and discern pathogenic versus non-pathogenic agents, or those which may become more replicative under environmental stress. Thus, a census of viral diversity will improve our future response when viruses impact the economic or ecological function of echinoderms.

Parvoviruses - linear, single-stranded DNA viruses - are the best documented group known to infect echinoderms since the discovery of a densovirus in a sea urchin metagenome from Hawaii in 2014 [19]. Shortly after, another densovirus was found in various sea star species and was implicated as the causative pathogen of the 2013/2014 Sea Star Wasting Syndrome (SSWS) outbreak in the Northeast Pacific [22]. However, subsequent attempts to correlate densoviruses to SSWS have not produced any clear association with pathology or disease [21–26]. Regardless, this discovery prompted a series of investigations into the diversity, prevalence, and association of these viruses with sea stars and SSWS [21, 24–26]. To date, no RNA virus identified using -omic approaches has been proven to cause any pathology in echinoderms [21, 27]. However, one clear line of evidence that has emerged from the accumulation of -omic data is that sea stars are infected by a diversity of viruses. We expect that they, and other echinoderms, will be host to a novel, undocumented diversity of RNA viruses.

Sequencing-based ‘viromics’ approaches have been the primary method for the discovery and characterization of echinoderm viruses. Other methods, like microscopy or culturing, are laborious and low throughput or hindered by the lack of available cell lines from aquatic invertebrates. All echinoderm virome studies, to date, have taken a viral metagenomic approach, where shotgun metagenomics is performed on encapsidated nucleic acids that have been enriched and selected for by chemical and/or nuclease treatment from size-filtered (<0.2 um) tissue homogenates [19–22, 25–27]. Metatranscriptomes and transcriptomes are increasingly used for viral discovery and have not yet been applied to echinoderms [28–31]. In the context of a metazoan, a metatranscriptome refers to the sequencing of total RNA (with rRNA removed) generally from samples pooled at the population level (multiple individuals of a species), while a transcriptome is the sequencing of poly-A tailed mRNA (with rRNA removed) generally from an individual organism [28, 32]. RNA-seq studies generating metatranscriptomes are typically for the purpose of viral discovery as opposed to transcriptomes, which are created for the purpose of analyzing gene expression patterns of an organism. In this study, we analyzed over 900 publicly available transcriptomes from echinoderms to characterize the biodiversity and distribution of RNA viruses. Together with previously published RNA viral metagenomes (i.e. a population of genomes of RNA viruses), we conducted a systematic survey of RNA viruses associated with the five major classes (Crinoidea, Holothuroidea, Asteroidea, Ophiuroidea and Echinoidea) of Echinodermata.

## Methods

The following sections detail the processing of the short-read libraries used for viral discovery and the analyses performed on the viral sequences. The two sources of libraries were transcriptomes and RNA-based viral metagenomes derived from various echinoderm species and tissues. All libraries processed in this study were obtained from the NCBI’s Short Read Archive (Table S1) and the assemblies generated from this study and the database used for viral discovery are accessible through the Open Science Foundation (https://osf.io/JXUAM). *Host transcriptomes and RNA-based viral metagenomes*

A total of 903 paired-end transcriptomes derived from the five classes of echinoderm hosts, including crinoid (n = 18; Crinoidea), sea cucumbers (n = 178; Holothuroidea), sea star (n = 179; Asteroidea), brittle star (n = 71; Ophiuroidea) and urchin (n = 457; Echinoidea). Raw sequences were quality controlled using Trimmomatic [33], to clip adapters, and FastX [34], to discard reads with lengths < 50 nt and average quality scores < 30. Transcriptomes were then assembled using default parameters in Trinity (v2.1.1) [35]. Contigs less than 500 nt were discarded prior to viral annotation.

A total of 24 paired-end RNA viral metagenomes derived from sea cucumbers (n=3; Holothuroidea) and sea stars (n=21; Asteroidea) were also retrieved from the Short Read Archive (Table S1). Raw sequences were merged, trimmed to clip adapters and discard reads with average quality scores < 20, and normalized to an average read depth of 100 using BBtools [36]. RNA viral metagenomes were assembled using Spades (v 3.11.1) with the -meta flag [37]. Contigs less than 500 nt were discarded prior to viral annotation.

### Virus discovery and annotation

We curated an RNA virus database for viral annotation. The database contained viral amino acid sequences from Shi et al 2016, Wu et al 2020 and Wolf et al 2020, and from the NCBI viral genome database after filtering for viral sequences from invertebrates and invertebrates/vertebrates [1, 29, 38]. Duplicated amino acid sequences were removed from the database using seqkit, yielding a total of 36,193 unique viral gene sequences. Echinoderm RNA viruses were then identified by querying transcriptome/RNA viral metagenome assemblies against the curated RNA viral database using DIAMOND BLASTx with the sensitivity parameter adjusted to ‘very-sensitive’ and an e-value cutoff of < 10^−20^ [39]. Contigs with significant similarity based on the BLAST criteria above were manually inspected in Geneious Prime (v 2020.2.2), and queried against the NCBI non-redundant database using the default BLASTp parameters to verify the viral annotation and assign putative gene function. BLASTp results were also used to identify conserved protein domains. Genome illustrations were created by exporting the sequence viewer from Geneious Prime (v 2020.2.2) into Adobe Illustrator (v25.4.1). Contigs with near identical matches to human and plant viruses and bacteriophage were removed. ORFs containing an RdRP domain were used for taxonomic placement. If a contig did not contain an RdRP sequence, a complete or partial ORF containing any conserved protein domain was chosen. Contigs that did not contain a conserved protein domain were removed from further analysis. Quality-filtered reads were mapped to viral contigs to obtain relative abundance information using BBMap [40] using the ‘semiperfect’ flag which accommodated ambiguous bases (with equivalent results achieved with the ‘perfect’ flag).

### Network analysis and phylogenetics

The taxonomic relationships of all recovered viral sequences were first mapped using a network analysis. To place sequences into broad taxonomic groups, we downloaded amino acid sequences from the top NCBI BLASTp results for each of our recovered viral sequences. A network was built based on sequence similarity using the online EFI-EST portal with default settings (minimum length = 0, maximum length = 50,000, filter type = e-value ≤ 10^−5^ [41]. Nodes represent individual viral sequences and edges are the degree of similarity based off BLASTp pairwise similarity scores using a minimum pairwise similarity of 35%. Clusters and singletons were removed from the network that did not contain any representative viral sequences with a RdRP sequence. The network was visualized in Cytoscape (v 3.8.2) using the ‘organic’ layout [42].

We further established the relatives of echinoderm RNA viruses based on RdRP phylogenies. Independent phylogenetic analyses were performed for viral orders using the type species designated by the International Committee on Taxonomy of Viruses. RdRP amino acid sequences were aligned using MAFFT [43] and phylogenies were inferred by a substitution model selected by smart model selection in PhyML 3.0 with branch support determined by bootstrapping for 100 iterations [44]. The resulting phylogenetic tree was visualized and annotated using FigTree v1.4.4 [45]. The *Amarillovirales* phylogeny was created from the MAFFT alignment used in [46].

## Results

We recovered a total of 347 viral contigs and 33 complete or near-complete genomes from the 927 short read libraries analyzed. A total of 259 viral contigs were recovered from transcriptomes, and 88 viral contigs from RNA viral metagenomes (Figure S1B). The mean viral contig length recovered from RNA viral metagenomes (mean ± standard deviation: 4,421 ± 3077 nt) was greater than from transcriptomes (3,143 ± 3102 nt; Figure S1A), and the size of sequencing libraries was weakly correlated with viral read depth in transcriptome libraries (p = 0.002, Pearson’s r = 0.29) but not in RNA viral metagenomic libraries (p = 0.82, r = 0.05; Figure S1C). On average the relative abundance of viral contigs as a proportion of total reads was low, (0.0085%), ranging from 0.000023% to 0.29% (x□= 0.0085%). The average percentage of viral reads in viral metagenomes (x□= 0.39%) was ∼45-fold higher than transcriptomes. However, viral abundance was highly uneven in the viral metagenomes with the average dropping to 0.07% (∼8-fold higher than transcriptomes) after excluding the top three most abundant samples.

Viral sequences were recovered from 111 of the 903 transcriptomic libraries and from all five echinoderm classes (Figure 1). Sea cucumbers exhibited the highest prevalence of viral contigs among all echinoderm libraries (*i*.*e*., individuals) screened, followed by sea urchins (10%, 45/457), and the highest proportion among transcriptomes screened (26%; 47/178) (Figure 1A). The majority of viral contigs recovered from transcriptomes came from adult tissue (70%) compared to embryos (20%) or larvae (10%) (Figure 1B). Transcriptomes derived from echinoderms during their larval stage had a slightly higher proportion of viral reads in their transcriptome (x□= 0.013%) than adults (0.0095%), and more than embryos (0.0034%; Figure 1C), but these differences were not significant (Kruskal-Wallis, chi-squared 1.65, p-value= 0.80).

**Figure 1:**
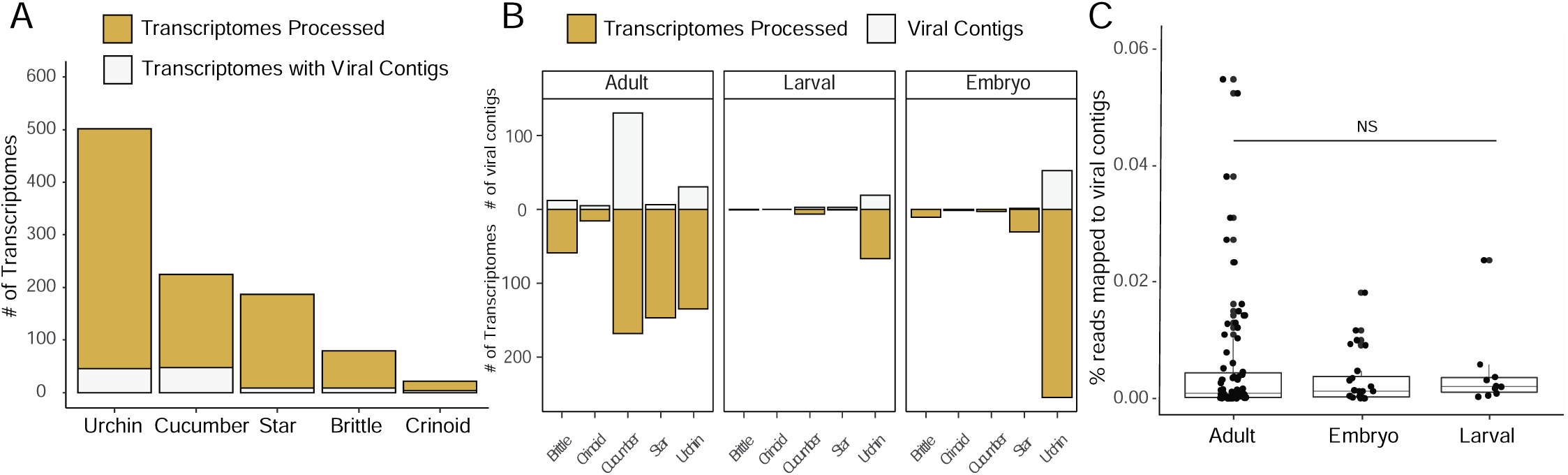
Summary of viral contigs discovered from echinoderm transcriptomes. (A) The number of transcriptomes downloaded from NCBI ordered by echinoderm class that were processed for viral discovery. (B) Top bars display the total number of viral contigs discovered separated by echinoderm class and life stage. Bottom bars display total number of transcriptomes separated by echinoderm class and life stage (C) Percentage of viral reads in transcriptomes by life stage. NS = non-significant.

Over half of the viral contigs contained an RdRP sequence (186/347), with 96 of these containing a complete or partial capsid sequence. Most viral contigs (215) contained at least a partial capsid sequence, and 42 viral contigs contained another conserved viral domain such as a methyltransferase or an RNA helicase domain (Supplemental Table 2). The majority of viral contigs were taxonomically placed in the order *Picornavirales* (n= 235) (Figure 2). The recovered picornaviruses were distributed among a variety of families with the largest number related to *Marnaviridae*, followed by *Dicistroviridae*, and *Iflaviridae* (Figure 3). Several unique clades within *Picornavirales* were represented by complete or near-complete genomes and may represent novel viral families (Figure 2). The second highest number of viral contigs were recovered from *Mononegavirales* (n = 12), with the remainder spread among seven orders (n= 20): *Durnavirales, Martellivirales, Nodamuvirales, Reovirales, Amarillovirales, Ghabrivirales*, and *Hepelivirales*. In general, the recovered viruses did not form new monophyletic clades within *Picornavirales* (Figure 3) or *Amarillovirales, Reovirales*, and *Mononegavirales* (Figure 4). However, within the *Hepivirales*, the recovered viruses formed a distinct monophyletic clade that is sister to the clade containing *Orthohepivirus* and *Piscihepivirus*.

**Figure 2:**
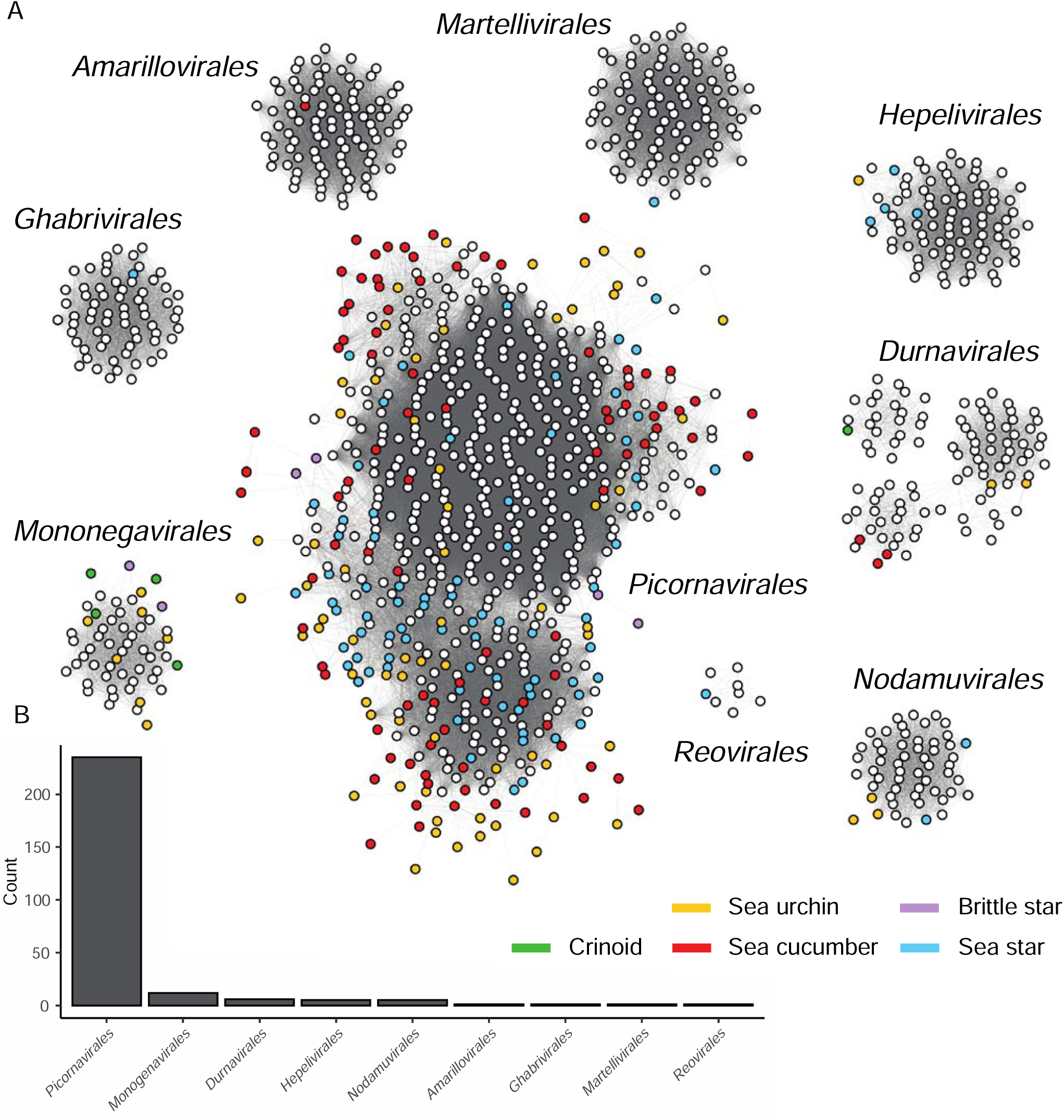
Picornaviruses are the dominant viral order found in echinoderms. (A) Colored circles represent viral sequences discovered from echinoderm transcriptomes and RNA viral metagenomes. White circles are viral genomes taken from NCBI. (B) The bar chart displays the number of echinoderm viruses discovered from each viral order.

**Figure 3:**
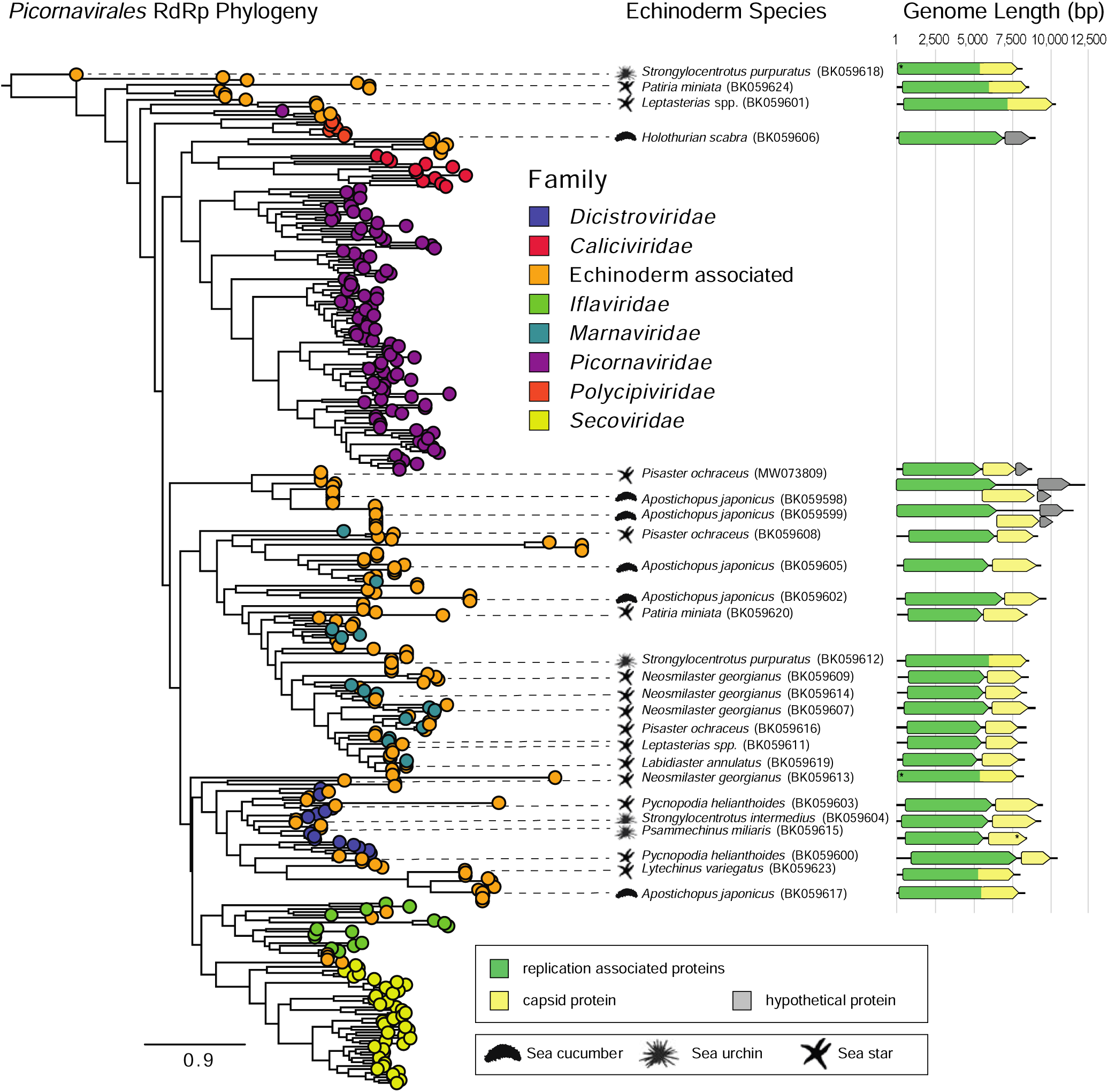
Echinoderm picornaviruses are broadly distributed across the *Picornavirales* phylogeny. Tips are colored by taxonomic family with orange circles representing echinoderm picornaviruses. Genome architectures of complete and near complete genomes recovered from assemblies displayed. Genomes are drawn approximate to scale in a 5’ to 3’ direction. Open reading frames denoted by boxes and colored by general function. Asterisk represents an incomplete open reading frame. Animal icons represent the echinoderm order the viral contig is associated with.

**Figure 4:**
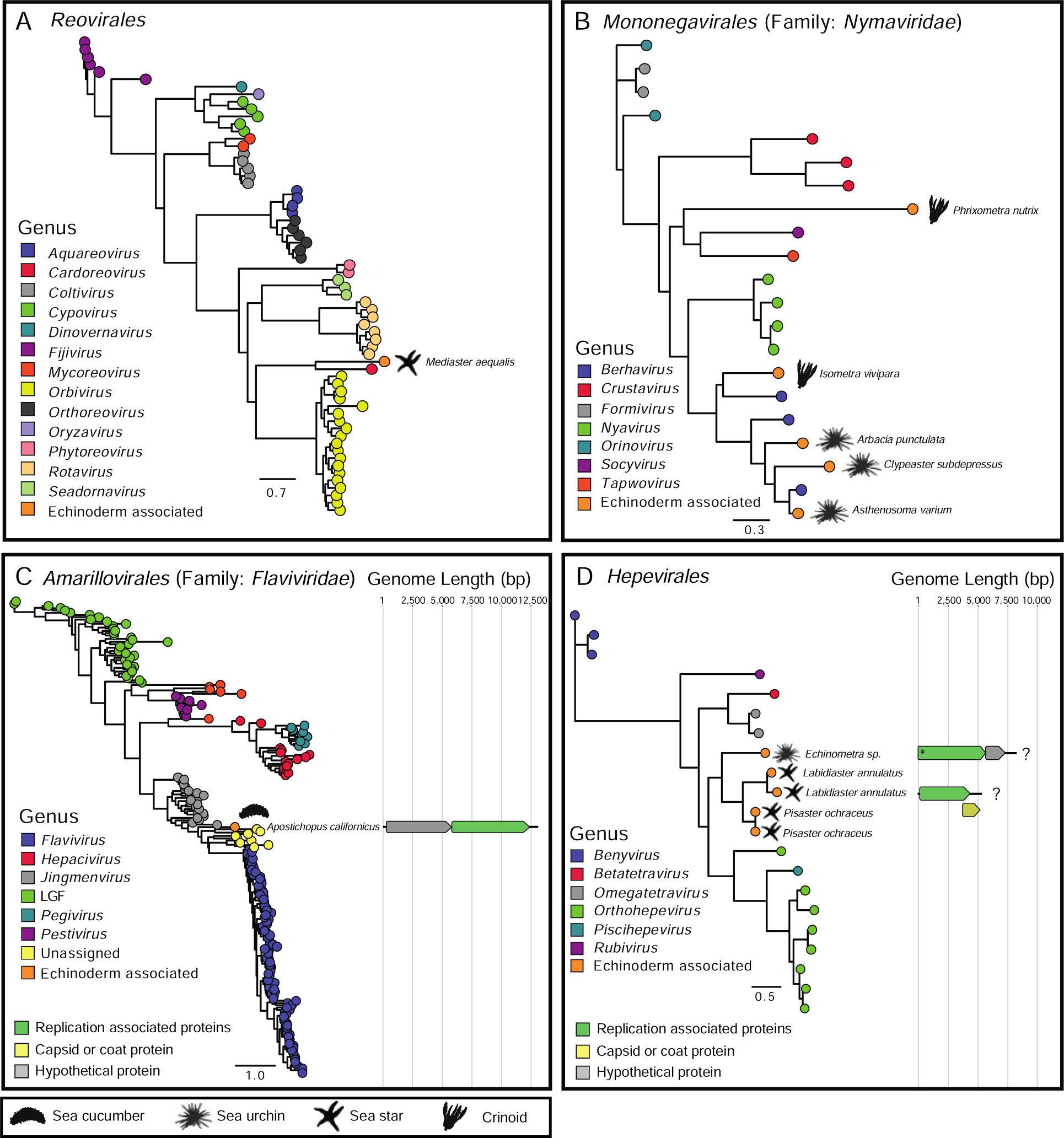
Phylogenetic placement of echinoderm viruses from *Reovirales, Mononegavirales, Amarillovirales*, and *Hepevirales*. Tips are colored by taxonomic family or genus with black circles representing echinoderm viruses. Genome architectures of complete and near complete genomes recovered from assemblies displayed. Genomes are drawn approximate to scale in a 5’ to 3’ direction. Open reading frames denoted by boxes and colored by general function. Asterisk represents an incomplete open reading frame and a question mark denotes potentially incompletegenome. Animal icons represent the echinoderm order the viral contig is associated with.

Viral contig lengths ranged from 502 nt to 12,989 nt. Complete and near complete genomes were recovered from *Picornavirales, Mononegavirales, Amarillovirales*, and *Hepelivirales* (Figure 5). The echinoderm picornavirus genomes exhibited three different open reading frame arrangements, which spatially separated the genome by function according to replication or encapsidation (Figure 3). The two most conserved protein domains related to replication were the RdRP (pfam00680) and RNA helicase (pfam00910) domains with many of the genomes also containing BIR (pfam00653), DSRM (pfam00035), Sigma70 (pfam04539), peptidases (pfam12381), and large tegument protein (PHA03247) domains. The conserved capsid domains found among the picornavirus genome included: rhv-like (pfam00073), dicistro VP4 (pfam11492), CRPV (pfam08762), and calici coat (pfam00915) domains (Figure 5). The recovered hepeviruses and mononegaviruses contigs contained the expected replication proteins but many lacked complete capsid proteins, indicating they were only near complete genomes (Figure 5). The flavivirus genome previously discovered in sea cucumber [27] was completed during our assembly, leading to the extension of a second complete ORF and extending the genome size from 8,883 to 12,989 nt.

**Figure 5:**
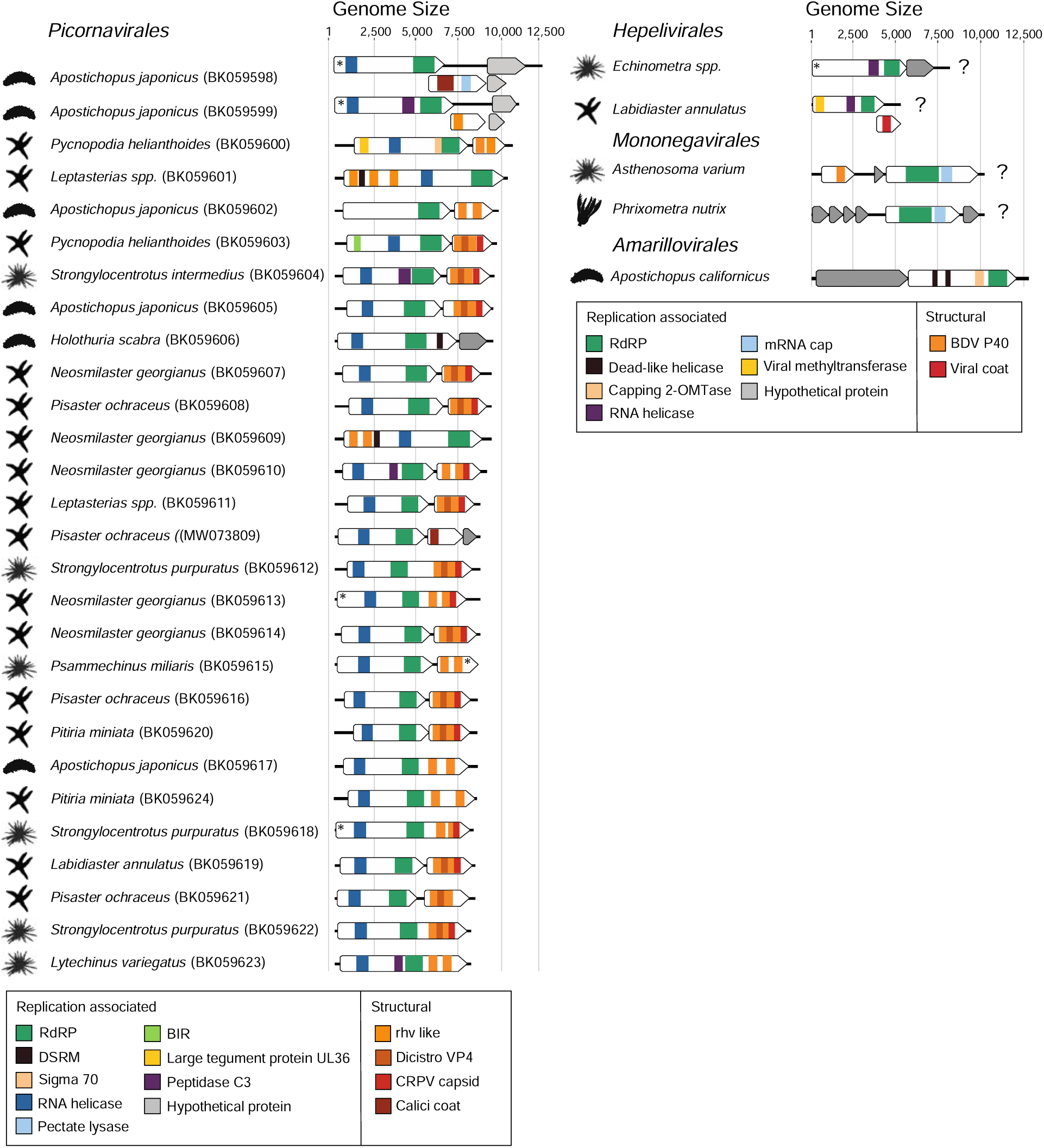
Genome architectures and comparison of complete or near complete genomes recovered from assemblies. Genomes are drawn approximate to scale in a 5’ to 3’ direction. Open reading frames denoted by boxes and colored regions represent conserved protein domains. Asterisk represents an incomplete open reading frame and question marks indicate missing open reading frames that would complete the genome. Animal icons represent the echinoderm order the viral contig is associated with.

## Discussion

The most prevalent RNA viruses in echinoderm transcriptomes and RNA viral metagenomes were picornaviruses which are non-enveloped, single-stranded RNA (+ssRNA) viruses. The order *Picornavirales* is comprised of eight families (*Caliciviridae, Dicistroviridae, Iflaviridae, Marnaviridae, Picornaviridae, Polycipiviridae, Secoviridae*, and *Solinviviridae*) and 103 genera and are among the most prevalent and diverse group of viruses found in -omics surveys of animal and environmental samples [47, 29, 38, 48]. The majority of the echinoderm-associated picornaviruses grouped into the *Dicistroviridae, Iflaviridae*, and *Marnaviridae* families, but other recovered viruses also comprised novel clades (Figure 3) which supported our expectation that the extant diversity of echinoderm RNA viruses is under sampled. The *Marnaviridae* are known to be ocean virioplankton which infect single-celled eukaryotes, such as phytoplankton and protists, but have also been found from metatranscriptomes from marine bivalves [48, 49]. It is possible that the *Marnaviridae* we observed infect protists that are symbionts or transiently associated with echinoderms. Alternatively, the host-range of the *Marnaviridae* family may extend beyond single-celled eukaryotes. The host range of many viral groups has changed considerably in recent years, and there are examples of host ranges within RNA viral families that do extend from protists to mammals, such as *Reoviridae* [50, 51]. All classified species of *Dicistroviridae* and *Iflaviridae* infect arthropods, and have largely been characterized due to the economic impacts of their pathogenicity though the host range of these families likely extends far beyond arthropods given their presence in metatranscriptomes from organisms in the phyla Mollusca, Cnidaria, and Platyhelminthes [31, 48, 52]. The disease severity from infections of both families ranges from inapparent to lethal, supporting the possibility that there may be non-pathogenic species infecting echinoderms.

Among the rare virosphere of echinoderms, and those that have few marine host associations, we observed *Martellivirales, Nodamuvirales, Reovirales, Amarillovirales, Mononegavirales*, and *Hepelivirales*. Many of these echinoderm viruses phylogenetically cluster with established invertebrate-infecting families or genera (i.e. *Nymaviridae* family within *Mononegavirales* or *Cardorevirus* genus within *Reovirales*) while some represent evolutionary novel lineages such as the flavivirus discovered from a sea cucumber (Figure 4C) [27]. Currently it is unclear if members of the rare virosphere infect all classes of echinoderms or if some are class specific. For example, the reovirus and flavivirus discovered in this study were only found in sea stars and sea cucumbers, respectively, though we cannot rule out methodological biases and insufficient sample size as proof of absence, requiring further research. Nevertheless, the discovery of these viruses significantly expands the host range for many of these groups and represents the first RNA viruses discovered from crinoids and brittle stars.

The vast majority of viruses recovered here, and elsewhere, using transcriptomic and metatranscriptomic approaches have been positive-sense single-stranded RNA (+ssRNA) [1, 48, 53]. This pattern of abundance likely has a basis in biology, but in the case of our dataset, may be inflated due to our use of transcriptomic data. The selection for polyadenylated transcripts during RNA-seq library preparation biases towards +ssRNA viruses, like picornaviruses, which have 3’polyadenylated tails [54, 55]. Studies utilizing a metatranscriptomic approach for RNA viral discovery generally do not find such a highly skewed distribution towards +ssRNA but are nevertheless the most abundant viral type [1, 29, 38, 48]. By utilizing viral RNA metagenomes and transcriptomes we have uncovered the fullest diversity of RNA viruses associated with echinoderms with the datasets available, though we expect future multi-omic efforts to reveal additional diversity.

The greatest difference between the two -omic approaches used in this study for viral discovery was the total number of viral contigs recovered and contig lengths. Viral RNA metagenomes generally contained > 3 viral contigs per library, which were ∼40 % longer, compared to transcriptomes, which contained 2.2 contigs per library. Additionally, the proportion of viral reads recovered exhibited a weak correlation with library size for transcriptomes but not for viral RNA metagenomes (Figure S1B). Thus, despite the efforts to enrich for viruses in the viral RNA metagenomes, the majority of libraries had a similar percent of viral reads compared to transcriptomes. Furthermore, these findings indicate that the efficacy of recovering RNA virus improves with sequencing depth, likely due to the improved assembly of sequenced found in low abundance.

The capacity for viral discovery using -omic approaches has greatly expanded our understanding of biodiversity and host range, fundamentally shifting the perception of viruses as solely pathogens to a more nuanced role as commensals or mutualists [1]. Performing a viral census of hosts, like echinoderms, provides a useful context about the prevalence and association of viruses that can help understand future outbreaks or changes in the susceptibility of marine animals due to stress from climate change and human activity. The full potential of -omic approaches to understand the biological or ecological role of the diversity of viruses uncovered will only be fully realized in partnership with advances in culturing techniques to study the infection of naïve specimens [56, 57]. Our study provides a comprehensive survey of RNA viruses present in echinoderm, contrasting the diversity and abundance of RNA viruses between echinoderm classes and life stages. We hope this information provides valuable context for advancing our understanding of the role of these viruses in marine hosts and ecosystems.

## Author statements

The authors declare that there are no conflicts of interest.

---

**Figure S1:**
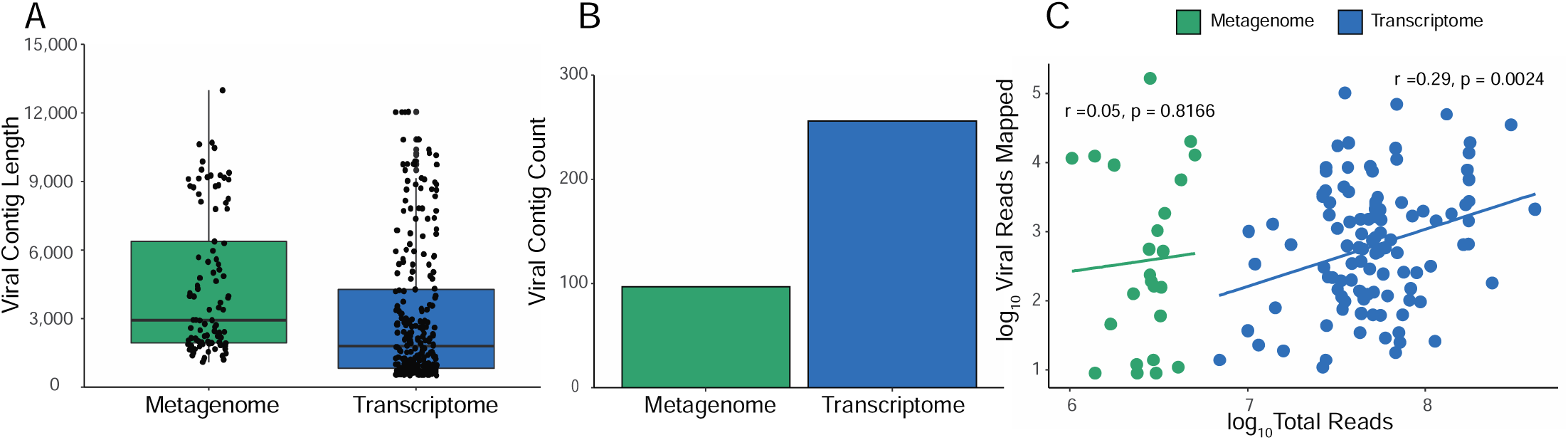
Summary statistics of viral discovery from short read libraries. (A) The distribution of viral contig lengths between RNA viral metagenomes and transcriptomes. (B) Total number of viral contigs discovered in RNA viral metagenomes and host transcriptomes.(C) Pearson’s correlations of total reads in a given library and total viral sequences in the same library.

